# Calcineurin is required for *Candida glabrata* Pdr1 transcriptional activation

**DOI:** 10.1101/2023.07.10.548434

**Authors:** Bao Gia Vu, Lucia Simonicova, W. Scott Moye-Rowley

**Affiliations:** Department of Molecular Physiology and Biophysics, Carver College of Medicine, University of Iowa, Iowa City, IA 52242, USA

## Abstract

Fluconazole is the most commonly used antifungal today. A result of this has been the inevitable selection of fluconazole resistant organisms. This is an especially acute problem in the pathogenic yeast *Candida glabrata*. Elevated minimal inhibitory concentrations (MICs) for fluconazole in *C. glabrata* are frequently associated with substitution mutations within the Zn2Cys6 zinc cluster-containing transcription factor-encoding gene *PDR1*. These mutant Pdr1 regulators drive constitutively high expression of target genes like *CDR1* that encodes an ATP-binding cassette transporter thought to act as a drug efflux pump. Exposure of *C. glabrata* to fluconazole induced expression of both Pdr1 and *CDR1*, although little is known of the molecular basis underlying the upstream signals that trigger Pdr1 activation. Here, we show that the protein phosphatase calcineurin is required for fluconazole-dependent induction of Pdr1 transcriptional regulation. Calcineurin catalytic activity is required for normal Pdr1 regulation and a hyperactive form of this phosphatase can increase resistance to the echinocandin caspofungin but does not show a similar elevation for fluconazole resistance. Loss of calcineurin from strains expressing two different gain-of-function forms of Pdr1 also caused a decrease in *CDR1* expression and fluconazole resistance, demonstrating that even these hyperactive Pdr1 regulatory mutants cannot bypass the requirement for calcineurin. Our data implicate calcineurin activity as a link tying azole and echinocandin resistance together via the control of transcription factor activity.

**Importance:** While drug resistant microorganisms are a problem in treatment of all infectious disease, this is an especially acute problem with fungi due to the existence of only 3 classes of antifungal drugs, including the azole drug fluconazole. In the pathogenic yeast *Candida glabrata*, mutant forms of a transcription factor called Pdr1 are commonly associated with fluconazole resistance and poor clinical outcomes. Here we identify a protein phosphatase called calcineurin that is required for fluconazole-dependent induction of Pdr1 transcriptional activation and associated drug resistance. Gain-of-function mutant forms of Pdr1 still required the presence of calcineurin to confer normally elevated fluconazole resistance. Previous studies showed that calcineurin is required for resistance to the echinocandin class of antifungal drugs and our data demonstrate this protein phosphatase is also required for azole drug resistance. Calcineurin plays a central role in resistance to two of the three major classes of antifungal drugs in *C. glabrata*.

## Results and discussion

Pdr1 is a central regulator of fluconazole resistance in *C. glabrata* and activates expression of the ATP-binding cassette transporter-encoding gene *CDR1* (reviewed in (1)). We have used a tandem affinity purification (TAP)-tagged form of Pdr1 to recover proteins that co-purify with this transcription factor. These data will be presented elsewhere but one of the proteins of interest that we recovered was the catalytic subunit of the protein phosphatase calcineurin (Cna1) (reviewed in (2)). Cna1 is well-established to control resistance to the echinocandin class of antifungal drugs via its activation of the Crz1 transcription factor (reviewed in (3)). Cna1 was also linked to fluconazole resistance as *cna1Δ* strains exhibited lowered azole drug resistance (4), although the underlying mechanism has remained unknown. Here we provide evidence that calcineurin is required for transcriptional activation by Pdr1.

To confirm the Pdr1:Cna1 association we detected by co-purification, we epitope tagged the *CNA1* gene with the insertion of two FLAG epitopes at the Cna1 carboxy-terminus to produce the Cna1-2X FLAG protein. This protein retained normal function and we prepared whole cell protein extracts from strains expressing wild-type or two different gain-of-function (GOF) forms of Pdr1. Pdr1 was immunoprecipitated from these strains and associated Cna1 was detected by immunoblotting using an anti-FLAG antibody (Supplementary Figure 1). Cna1 was found to associate with all the forms of Pdr1 we examined. These data confirm that Cna1 is associated with Pdr1 in vivo, although we do not yet know if this represents a direct or indirect association.

Having established that Cna1 was associated with Pdr1 in vivo, we analyzed the effect of Cna1 on Pdr1-dependent transcriptional activation by comparing isogenic wild-type and *cna1Δ* cells. We reproduced the published phenotypes caused by loss of Cna1 as a *cna1Δ* strain exhibited a pronounced increase in susceptibility to both caspofungin and fluconazole (Figure 1A). Importantly, the normal fluconazole-induced transcriptional activation of both *PDR1* and *CDR1* genes was lost in the *cna1Δ* strain (Figure 1B). These data argue that the effect of Cna1 on fluconazole resistance comes about by stimulating function of the Pdr1 transcription factor. Expression of the gene (*ERG11*) encoding the fluconazole target enzyme were unaffected by the loss of calcineurin activity. The impact of calcineurin on expression of Cdr1, Pdr1 and Erg11 was also seen by western blot analysis using antibodies directed against each of these proteins (Supplementary Figure 2A and 2C). Normal calcineurin activity is required for wild-type and fluconazole-induced expression of *CDR1* and *PDR1* but dispensable for *ERG11*.

**Figure 1.**
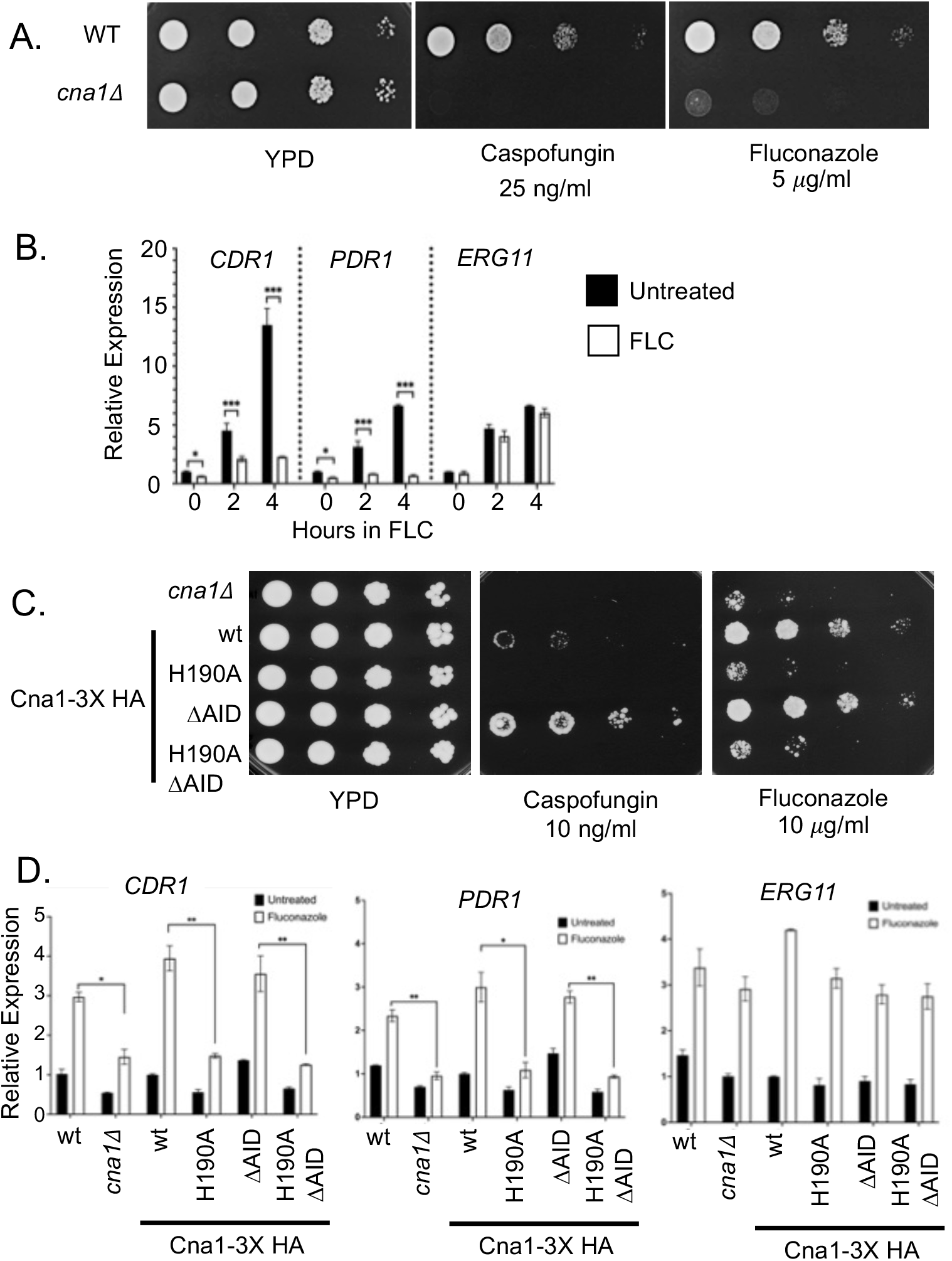
Calcineurin is required for fluconazole-induced *CDR1* and *PDR1* expression. A. Isogenic wild-type and *cna1Δ* strains were grown to mid-log phase and serially diluted on YPD medium containing the indicated concentrations of caspofungin and fluconazole. B. Wild-type and *cna1Δ* strains were grown to mid-log phase and then treated with 20 μg/ml fluconazole for the indicated times on the X axis. C. A strain lacking the *CNA1* gene (*cna1Δ*) was transformed with a low-copy-number plasmid expressing the indicated forms of Cna1 as 3X HA-tagged proteins: the wild-type protein (wt), a catalytically inactive mutant protein (H190A), a constitutively active phosphatase mutant (Δ AID) or a double mutant (H190A Δ AID). Transformants were grown to mid-log phase and serial dilutions spotted on plates containing the indicated medias. D. The transformants expressing the indicated forms of Cna1, along with wild-type and isogenic *cna1Δ* control strains, were grown to mid-log phase, left untreated or challenged with fluconazole as above and then assayed for levels of *CDR1, PDR1* and *ERG11* mRNA using RT-qPCR.

We took advantage of the extensive characterization of Cna1 (discussed in (2)) and tested two different mutant forms of this factor to assess the effect on Pdr1 transcriptional activation. We prepared a catalytically inactive mutant derivative (H190A Cna1) and a constitutively active form of Cna1 that lacks the autoregulatory inhibitory domain (AID) that would normally restrain phosphatase activity (reviewed in (5)). These mutations were introduced into a *cna1Δ* strain and tested for the ability to complement the caspofungin and fluconazole hypersusceptibility of this strain as well as to correct the defective fluconazole induction of *PDR1* and *CDR1* transcription.

The presence of the Δ AID Cna1 protein led to elevated caspofungin resistance as has been seen for ion stress (6) but only restored fluconazole susceptibility to normal levels (Figure 1C). To our knowledge, the elevated caspofungin resistance of the Δ AID Cna1 protein has not been reported before. Loss of the catalytic histidine residue from both forms of Cna1 led to loss of complementing activity. While calcineurin activity is limiting for resistance to caspofungin, only normal levels are required for wild-type fluconazole susceptibility. This is supported by levels of fluconazole-inducible transcription of both *CDR1* and *PDR1* as loss of the catalytic histidine blocked the drug-inducible expression of both these genes and the loss of the AID restored normal levels of expression (Figure 1D). Note that the effects of calcineurin are restricted to the Pdr1/*CDR1* axis of fluconazole resistance as the response of the *ERG11* gene to fluconazole was unaffected by alterations in calcineurin activity.

Calcineurin activity requires both a catalytic (*CNA1*) and a regulatory subunit (*CNB1*). To ensure the regulatory subunit was also involved in the control of *CDR1* and *PDR1* expression, we prepared a *cnb1Δ* strain. To confirm the specificity of the effect of loss of calcineurin activity, we also generated a strain lacking the *PPT1* gene that encodes a Cna1-related protein serine/threonine phosphatase (7). Loss of *CNB1* led to an increase in fluconazole susceptibility similar to that caused by the *cna1Δ* allele (Figure 1A) while loss of *PPT1* had no significant effect on this phenotype (Supplementary Figure 3A). We analyzed Cdr1, Pdr1 and Erg11 expression by western blotting (Supplementary Figure 3B and 3C). As the fluconazole phenotype suggested, the *cnb1Δ* strain did not induce either Cdr1 or Pdr1 while fluconazole induction was normal for the *ppt1Δ* strain. Erg11 expression was unaffected by loss of either gene.

We also probed the effect of the immunosuppressive drug FK506 which is well-established as a potent inhibitor of calcineurin activity (8). Addition of 1 μM of FK506 led to a striking reduction in the level of resistance to both fluconazole and caspofungin in wild-type cells (Supplementary Figure 4A). Importantly, the presence of FK506 completely blocked the fluconazole-dependent induction of *PDR1* transcription (Supplementary Figure 4B) as well as the increased levels of Pdr1 protein (Supplementary Figure 4C and 4D) while no significant effect was seen on the expression of *ERG11*.

Having established that calcineurin was required for both wild-type and fluconazole-induced levels of Pdr1 function, we tested the role of this phosphatase on two different gain-of-function alleles of *PDR1*. We disrupted the *CNA1* gene in three isogenic backgrounds containing the wild-type, D1082G or R376W forms of *PDR1*. First, we tested these strains for their fluconazole susceptibility. Loss of Cna1 from either GOF-expressing form of Pdr1 caused an increase in azole susceptibility although considerable resistance was maintained (Figure 2A). We also assessed the effect of a *cna1Δ* allele on the level of wild-type and fluconazole-induced *CDR1* and *PDR1* expression at both the RNA and protein level (Figure 2B, 2C, 2D and 2F). In all cases, loss of Cna1 reduced expression of both *CDR1* and *PDR1*. This reduction correlated well with the observed increase in fluconazole susceptibility of these GOF strains lacking calcineurin and argues that calcineurin activity is required for full function of even these GOF forms of Pdr1.

**Figure 2.**
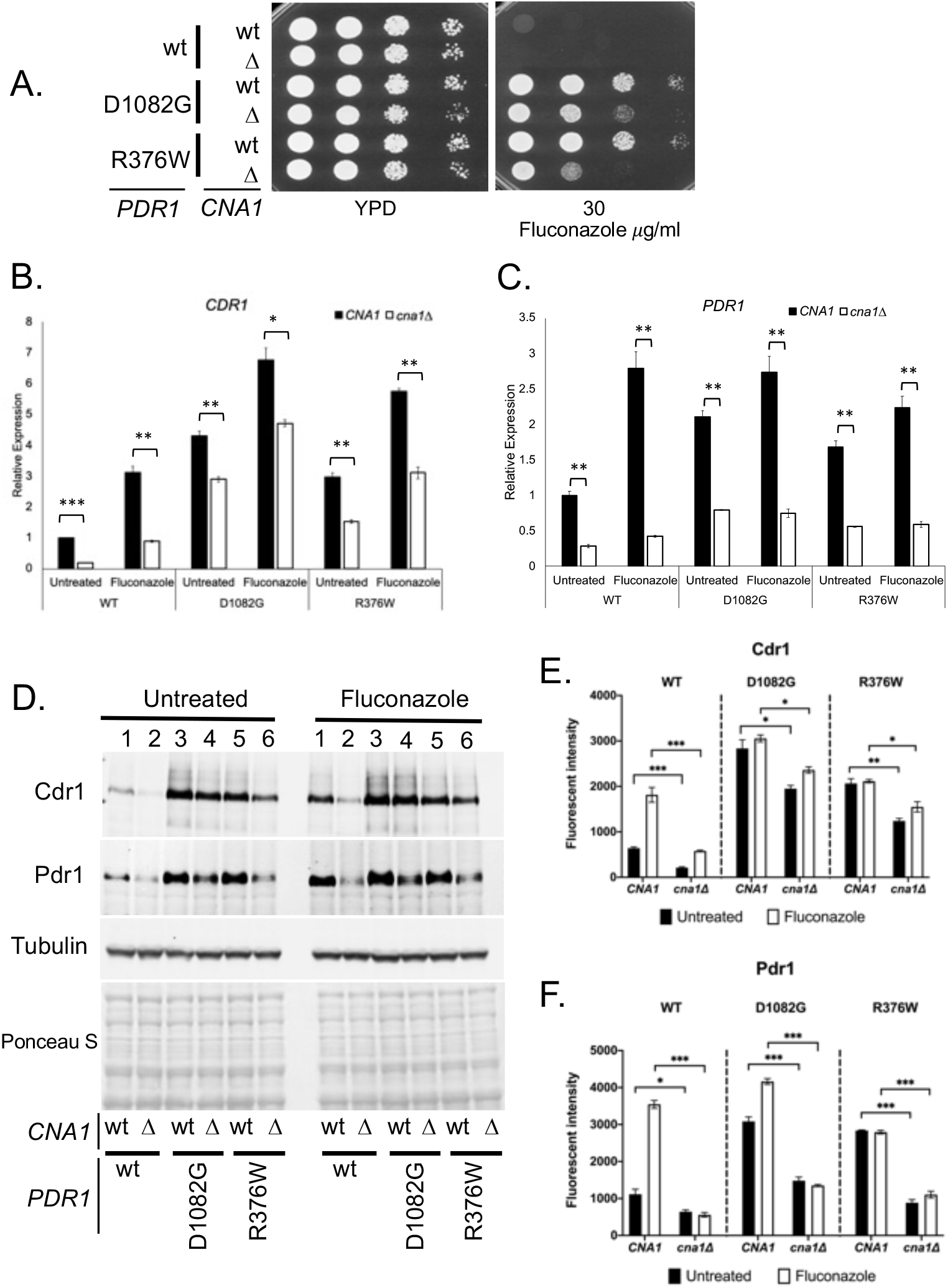
Calcineurin is required for normal activity of gain-of-function forms of Pdr1. A. Isogenic strains containing or lacking the *CNA1* gene and expressing the indicated forms of Pdr1 were grown to be tested for resistance to elevated level of fluconazole indicated as described before. B. The strains listed above were tested for the level of untreated and fluconazole-inducible transcription of *CDR1* and *PDR1* by RT-qPCR as above. C. Whole cell protein extracts were prepared from the strains treated as in B and analyzed by western blotting using polyclonal antibodies directed against Cdr1 or Pdr1 and a mouse monoclonal that detects tubulin protein to ensure equal loading. The transferred proteins were also stained by Ponceau S after western blotting to confirm equivalent transfer and protein extraction. Quantitation of the western blot signals for Cdr1 (E) and Pdr1 (F) is also shown in the respective panels.

These findings identify calcineurin as a critical upstream modulator of susceptibility to two of the most important classes of antifungal drugs: azoles and echinocandins. The mechanism of calcineurin activation of Crz1 for echinocandin resistance (3) and Pdr1 for fluconazole resistance seems to be different, at least genetically, since a hyperactive allele of *CNA1* leads to echinocandin resistance that is higher than that seen in the wild-type strain while while this same allele simply restores azole susceptibility to normal (Figure 1 and supplementary Figure 2). We also provide an explanation for the previous observed increase in fluconazole susceptibility of a *cna1Δ* strain (4) as this strain exhibits a defect in the expression of both *CDR1* and *PDR1* under wild-type and fluconazole-challenged conditions. Calcineurin is also required for full function of two different GOF alleles of *PDR1*, a finding that indicates the increased activity of these transcription factors must involve more than a single positive input.

The GOF forms of Pdr1 maintain significant expression of both Pdr1 itself and also the *CDR1* target gene in the absence of Cna1. While we see association of Cna1 with Pdr1 in co-immunoprecipitation assays, we do not yet know if this interaction is direct. The mechanism underlying control of Pdr1 function by calcineurin catalytic activity is a main focus of future experiments. Our preliminary evidence suggests that bulk phosphorylation of Pdr1 is increased upon fluconazole challenge (data not shown) which suggests that the direct calcineurin target may be another factor but this remains to be determined. Calcineurin is known to be stimulated by elevated Ca^2+^ levels in the cell and has been previously shown to impact fluconazole susceptibility (4). Fluconazole is fungistatic on *C. glabrata* but can be converted to fungicidal if combined with calcineurin inhibitors (9). We speculate that part of this conversion may involve inhibition of the calcineurin-dependent activation of the Pdr1/*CDR1* fluconazole resistance axis we describe here.

## Supplementary Figures and Material and Methods

**Supplementary Figure 1.**
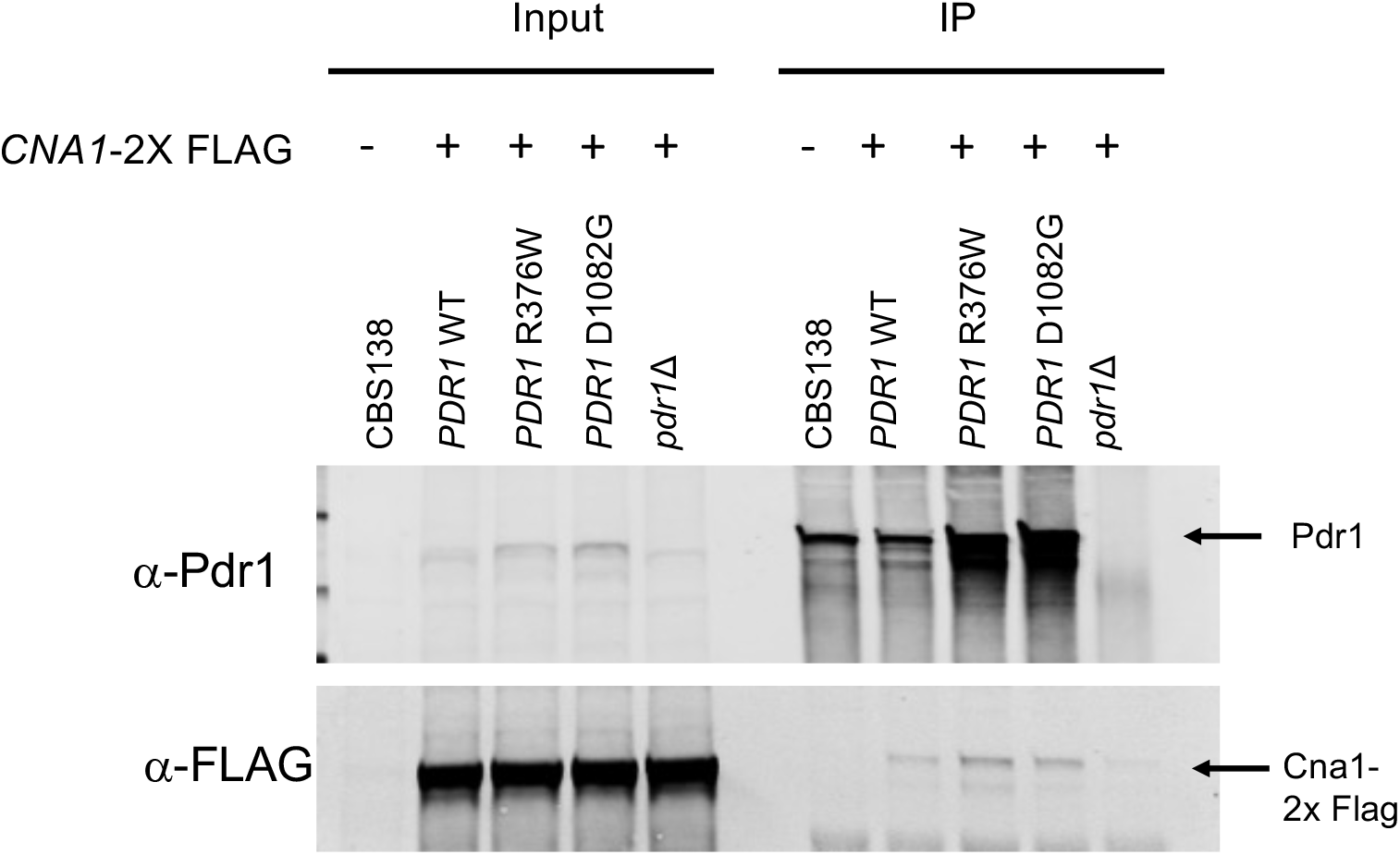
Co-immunoprecipitation of Pdr1 with epitope-tagged Cna1. Transformants expressing various forms of *PDR1* and the low-copy-number *CNA1*-2X FLAG-tagged allele were grown to mid-log phase, treated with 20 mg/ml of fluconazole for 2 hours at 30°C. Whole cell protein extracts were then prepared under nondenaturing conditions and subjected to immunoprecipitation using anti-Pdr1 antibody to recover Pdr1 and associated proteins. A fraction (1%) of the total extract was reserved to serve as an input control (Input) with the remainder used for anti-Pdr1 immunoprecipitation. Immunoprecipitates were washed and then separated on SDS-PAGE followed by western blotting analysis using polyclonal anti-Pdr1 and mouse monoclonal anti-FLAG antisera. Location of the proteins of interest is indicated by the arrows.

**Supplementary Figure 2.**
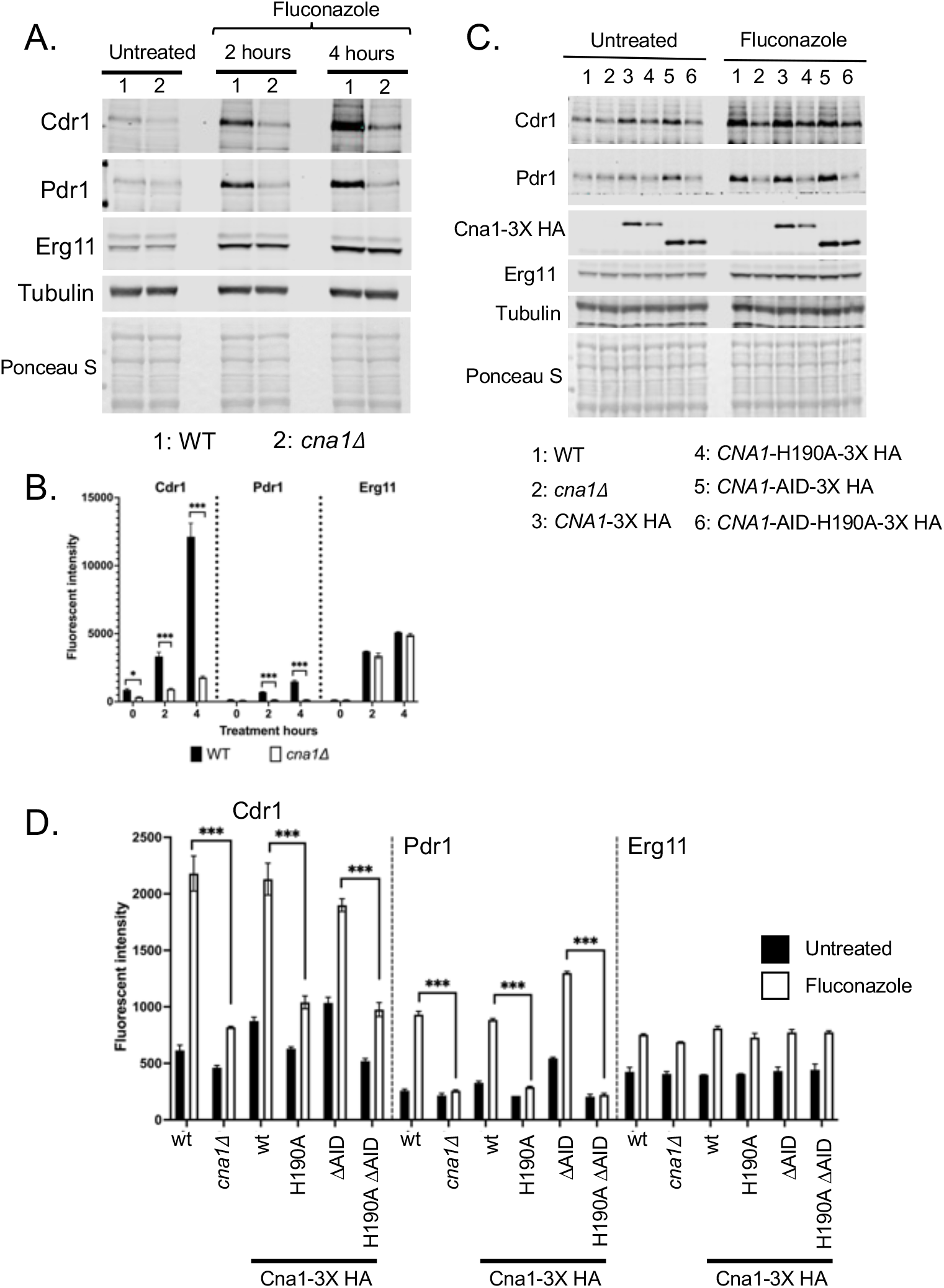
Western blot analyses of the role of Cna1 in expression of Pdr1 target genes. A. Isogenic wild-type and *cna1Δ* strains were grown to mid-log phase and then challenged with 20 μg/ml fluconazole for the times indicated as described in Figure 1A and 1B. Whole cell protein extracts were prepared under denaturing conditions were prepared and analyzed by western blotting using polyclonal antisera detecting Cdr1 or Pdr1, an anti-peptide antibody detecting Erg11 or a mouse monoclonal against tubulin. The transferred proteins were stained with Ponceau S on the membrane to ensure equivalent transfer and loading. B. Quantitation of the western blotting in part A is shown. C. Western blotting of different proteins of interest as described in Figure 1C and 1D. Note the equivalent levels of the Cna1-2X HA proteins indicating that the defects caused by the catalytic mutant form of Cna1 (H190A) are not due to lack of expression of proteins containing this mutation. D. Quantitation of the expression of the indicated proteins in panel C is shown.

**Supplementary Figure 3.**
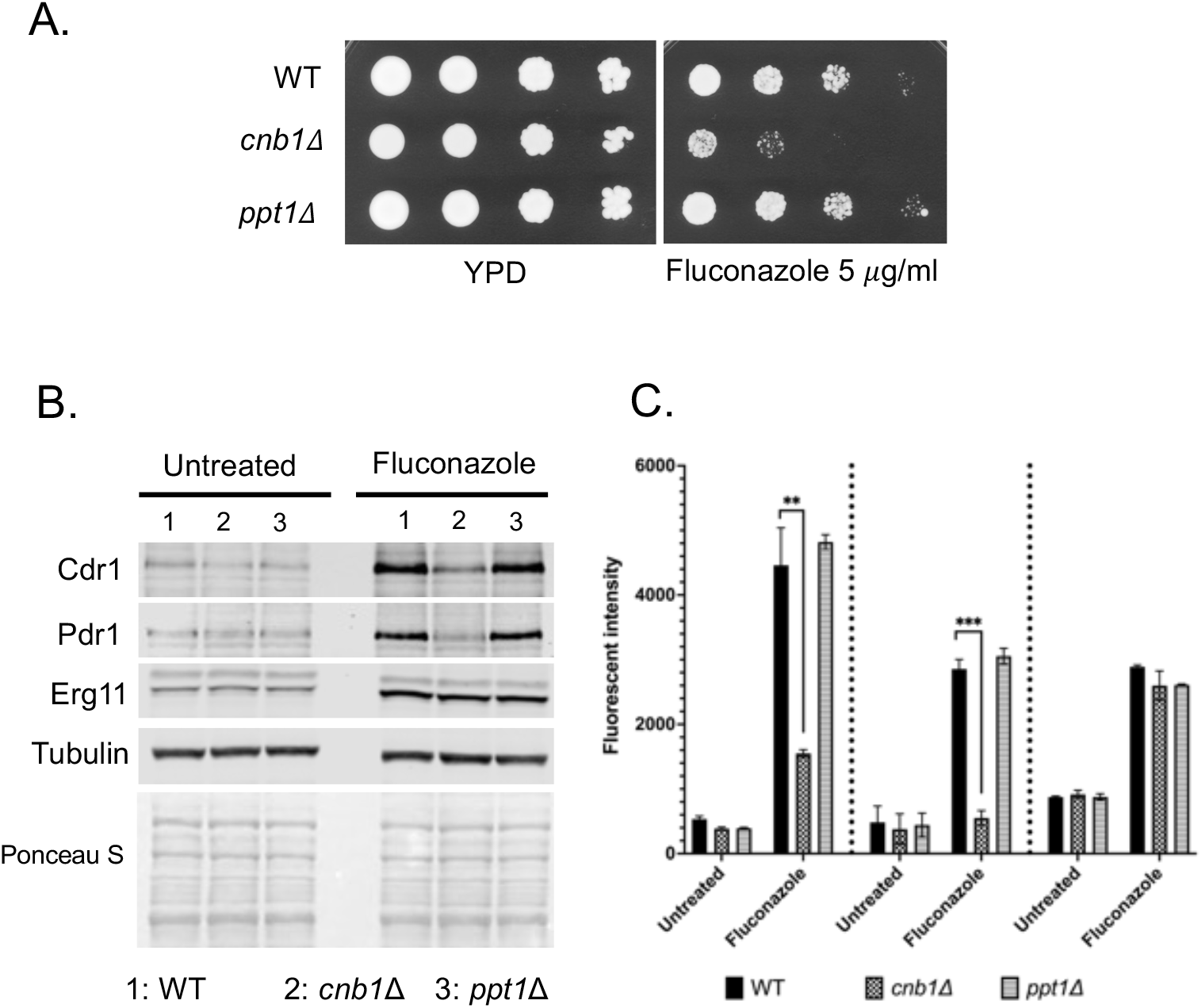
Defective Pdr1 regulation is reproduced by loss of the calcineurin regulatory subunit but not a different protein phosphatase. A. Isogenic strains lacking either the calcineurin regulatory subunit-encoding gene (*CNB1*) or a different serine/threonine protein phosphatase (*PPT1*) were tested for resistance to the indicated concentration of fluconazole as described previously. B. Expression of the indicated proteins was evaluated by western blotting as above. C. Quantitation of the western blot assay in B is shown.

**Supplementary Figure 4.**
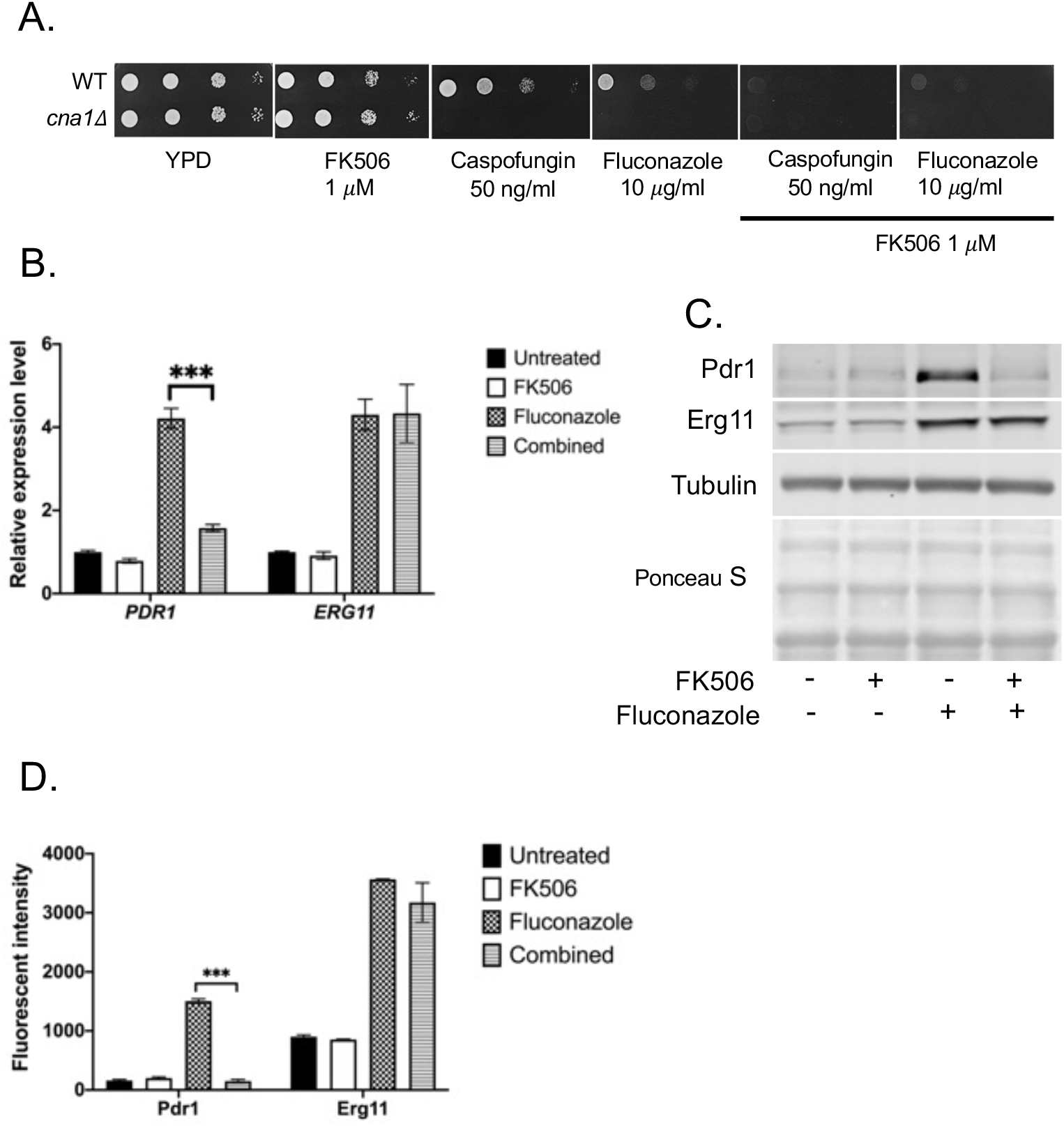
Chemical inhibition of calcineurin activity phenocopies genetic loss of the phosphatase. A. Isogenic wild-type and *cna1Δ* strains were analyzed by serial dilution on YPD medium containing the indicated drugs. FK506 is an inhibitor of calcineurin function and was added alone or in combination with caspofungin or fluconazole. B. Wild-type cells were grown to mid-log phase and then treated with FK506, fluconazole or both drugs. After two hours, *PDR1* and *ERG11* mRNA levels were assessed by RT-qPCR assay. C. Western blot analysis of cells treated as above using the indicated antisera. D. Quantitation of the western blot data in C.

## Materials and methods

### Reagents, *C. glabrata* strains and growth conditions

Fluconazole was purchased from LKT (laboratories, St Paul, MN). Caspofungin was purchased from Apexbio (Houston, TX). FK506 was purchased from Enzo Life Sciences (Farmingdale, NY). General growth conditions, nourseothricin section, and recyclable marker eviction techniques were previously described in (10). All strains used in this study are listed in the supplemental table 1. Resistance phenotypes were assayed by plating serial dilutions of log phase cultures on solid media. Transformations were carried out using a standard lithium acetate protocol.

### Strain and plasmid construction

*CNA1, CNB1*, and *PPT1* gene deletion constructs were made by assembling the recyclable cassette from pBV65 (10) and fragments from the immediate upstream/ downstream regions of each gene by Gibson assembly cloning (New England Biolabs, Ipswich, MA). Eviction of the recyclable cassette left a single copy of *loxP* in place of the removed target gene coding region. Standard homologous transformation was used and all disruption mutations were verified by PCR using appropriate primers.

The 3X HA epitope was inserted into the C-terminus of the *CNA1* gene in place of the native stop codon. The *CNA1-*3X HA cassette was cloned into the pBV133 vector (11), which carries a nourseothricin selection maker. From this plasmid, carrying the wildtype version of *CNA1*-3X HA, *CNA1*-H190A-3X HA, *CNA1*-AID-3X HA, and *CNA1*-AID-H190A-3X HA plasmids were subsequently generated from Gibson assembly cloning (New England Biolabs).

To clone the *CNA1* allele as a C-terminal fusion with a 2X FLAG tag, the low-copy-number vector pCnat-*CNA1*-3X HA described above was linearized with the restriction enzymes SacI and KpnI to replace the 3X HA tag with 2X FLAG. The 2X FLAG sequence was introduced via PCR as an oligonucleotide primer along with a reverse primer that produced the 3’ *CNA1* UTR and cloned by use of the Gibson Assembly Cloning Kit (NEB #E5510S). The identity of the vector pCnat-*CNA1*-2X FLAG was also confirmed by sequencing.

### Quantification of transcript levels by RT-qPCR, antibodies and western immunoblot analysis

Detailed protocols were previously described (11-13). Target gene transcript levels were normalized to transcript levels of 18S rRNA. The HA antibody (clone 2-2.2.14) was purchased from Invitrogen (Carlsbad, CA). The Pdr1, Cdr1, and Erg11 antibodies were previously described (12-14).

### Statistics

The Student T-test was used to assess the statistical significance of results of comparisons of samples. Paired conditions were used for comparisons of results from the same isolate obtained under different treatment conditions, while unpaired conditions were used for comparisons of results from isolates obtained under the same treatment conditions (*, P < 0.5; **, P < 0.01; ***, P < 0.001).

**Supplemental table 1:** Strain list.

| Strain name | Parent | Genotype | Reference |
| --- | --- | --- | --- |
| CBS138 (ATCC2001) | CBS13<br>(ATCC2001) | Wildtype isolate | ATCC |
| BVGC65 | CBS138 | *pdr1Δ::loxP* | (10) |
| BVGC379 | CBS138 | *cna1Δ::loxP* | This study |
| BVGC600 | CBS138 | *cnb1Δ::loxP* | This study |
| BVGC612 | CBS138 | *ppt1Δ::loxP* | This study |
| BVGC84 | BVGC65 | *PDR1::loxP* | (11) |
| BVGC86 | BVGC65 | D1082G *PDR1::loxP* | (11) |
| BVGC87 | BVGC65 | R376W *PDR1::loxP* | (11) |
| BVGC390 | BVGC84 | *PDR1::loxP cna1Δ::loxP* | This study |
| BVGC394 | BVGC86 | D1082G *PDR1::loxP*<br>*cna1Δ::loxP* | This study |
| BVGC397 | BVGC87 | R376W *PDR1::loxP*<br>*cna1Δ::loxP* | This study |

